# Structural insights into unique features of the human mitochondrial ribosome recycling

**DOI:** 10.1101/399345

**Authors:** Ravi K. Koripella, Manjuli R. Sharma, Paul Risteff, Pooja Keshavan, Rajendra K. Agrawal

## Abstract

Mammalian mitochondrial ribosomes (mitoribosomes) are responsible for synthesizing proteins that are essential for oxidative phosphorylation or energy (ATP) generation. Despite their proposed bacterial origin, the composition and structure of the human mitoribosome and its translational factors are dramatically different from their bacterial counterparts. The mammalian mitoribosome recycling factor (RRF_mt_) carries a mito-specific N-terminus extension (NTE), which is necessary for the function of RRF_mt_. Here we present a 3.7 Å resolution cryo-EM structure of the human 55S mitoribosome-RRF_mt_ complex, which reveals α-helix and loop structures for a portion of the NTE that makes multiple mito-specific interactions with functionally critical regions of the mitoribosome. These include ribosomal RNAs segments that constitute the peptidyl transferase center (PTC), those that connect PTC with the GTPase-associated center, and with multiple mitoribosomal proteins. Our structure also reveals novel conformational changes in mitoribosome due to RRF_mt_ binding. Together, these findings help understand the unique features of mitoribosome recycling.

## Introduction

Ribosomes are highly complex macromolecular structures that orchestrate the process of protein synthesis in coordination with messenger RNAs (mRNAs), transfer RNAs (tRNAs) and multiple translational factors. Mitochondria are cellular organelles that carry their own genetic material and gene-expression machinery including their own ribosomes. Mammalian mitochondria synthesize 13 polypeptides that form essential components of the oxidative phosphorylation machinery ^1^. Although the ribosomal RNA (rRNA) components of the mitochondrial ribosome (mitoribosome) are encoded by its own mito (mt)-DNA, all the proteins required for mitochondrial translation including the mt-ribosomal proteins and translation factors are encoded by nuclear genes, translated by the cytoplasmic ribosomes and then transported into the mitochondria ^2^. The mitoribosomes and their associated translation machinery are distinct from those in the cytoplasm and display features reminiscent of prokaryotic translation ^3^, in line with the assumption that mitochondria have evolved from endocytosis of an α-proteobacterium by an ancestral eukaryotic cell ^3^.

The basic features of ribosome recycling are known mostly through biochemical and structural studies from bacterial system. After the termination step of protein synthesis, the mRNA and a deacylated tRNA in its P/E state remain associated with the ribosome ^4,5^, and the complex is widely referred to as the post termination complex (PoTC). To initiate a new round of protein synthesis, the ribosome-bound ligands must be removed from the PoTC and the ribosome must be split into its two subunits. Disassembly of the PoTC requires a concerted action of two important protein factors, the ribosome recycling factor (RRF) and the elongation factor G (EF-G) ^4,6,7^. The binding of EF-G in conjugation with guanosine 5′-triphosphate (GTP) to the RRF-bound PoTC dissociates the 70S ribosome into its two subunits upon GTP hydrolysis ^8^.

High-resolution crystallographic studies of RRF from several bacterial species revealed its two-domain structure that adopts nearly an “L” conformation ^9,10^. While the larger domain I is composed of three long α-helical bundles, the smaller domain II is an insert with α/β motif. The two structural domains are held together by linker regions that confer conformational flexibility essential for the proper functioning of the molecule. Previous hydroxyl radical probing experiments ^11^, cryo-electron microscopic (cryo-EM) reconstructions ^12^-^15^, and X-ray crystallographic structures ^16^-^18^ of RRF-bound ribosome and functional PoTCs have revealed that, while domain I of RRF occupies an almost identical binding position in various RRF-bound 70S structures, domain II has been found to be oriented differently in relation to domain I, and thus inferred to be the major structural component driving RRF-mediated ribosome recycling ^19^. Domain I binds to the 50S subunit, spanning over its binding sites for peptidyl (P) and aminoacyl (A) tRNAs and interacting with some key elements of the 23S rRNA, including the P-loop of the peptidyl transferase center (PTC) ^20^, helices 69 (H69) and H71, whereas domain II localizes to the intersubunit space in close proximity to the protein L11 stalk-base from the large subunit and protein S12 from the 30S subunit ^12-15,17^. Cryo-EM studies of the spinach chloroplast ribosome, in complex with chloroplast-RRF (chl-RRF), which carries an extended N-terminus, and hibernation-promoting factor PSRP1 have shown that the overall binding position of chl-RRF on chlororibosome is similar to that in bacterial ribosome ^21,22^. However, structure of the extended N-terminus of the chl-RRF is unknown. The interaction of RRF with helices H69 and H71, which are involved in the formation of two of the strongest and highly conserved inter-subunit bridges B2a and B3, respectively, had led to the interpretation that RRF-induced disruption of the inter-subunit bridges facilitate subunit separation ^12-14,16,17^.

Despite anticipated similarity between the bacterial and mitochondrial ribosomes, the first mitoribosome cryo-EM structure had revealed that the mammalian mitoribosomes have diverged considerably from their bacterial counterparts and acquired several unique features ^23^, a finding that was subsequently confirmed in higher-resolution studies ^24^-^26^. The most striking difference is the reversal in the protein to RNA ratio; whereas the bacterial ribosomes are high in rRNA the mitoribosomes are high in protein ^27^. Despite significant increase in protein mass due to acquisition of mitochondrial-specific proteins and addition of insertions and extensions to many homologous ribosomal proteins ^23^-^26^, the overall sedimentation coefficient of the mitoribosome is 55S, and those of its small and large ribosomal subunits are 28S and 39S, respectively. Furthermore, most mitochondrial translational factors have also acquired insertions and extensions as compared to their bacterial counterparts ^28,29^. Thus, even though the mitochondrial translation closely resembles the prokaryotic system in terms of overall sequence of events and the accessory protein factors involved, there are considerable differences as well. For example, only two initiation factors, IF2_mt_ and IF3_mt_ have been identified in mammalian mitochondria and homologue for IF1 is absent ^30^-^32^. Unlike in bacteria, where a single EF-G molecule participates in both the elongation and ribosome recycling steps ^33^, mammalian mitochondria utilizes two isoforms of EF-G, EF-G1_mt_, which functions as a translocase during the polypeptide elongation step ^34^, and EF-G2_mt_ ^35^, which works exclusively with RRF_mt_ ^36^ to catalyze the mitoribosome recycling.

The amino acid (aa) sequence of the human RRF_mt_ is about 25-30 % identical to its bacterial homologues and carries an additional 80 aa long extension at its N-terminus ^37^. RRF_mt_ was shown to be essential for the viability of human cell lines and depletion of this factor leads to mitoribosomal aggregation and mitochondrial dysmorphism ^38^. Here we present a 3.7-3.9 Å resolution cryo-EM structure of human RRF_mt_ in complex with the human 55S mitoribosome, and investigate the molecular interactions of RRF_mt_ and its mito-specific N-terminal extension (NTE) with the 55S mitoribosome to gain insights into the process of mitoribosome recycling.

## Results and Discussion

### RRF_mt_ binds to the model mitoribosome PoTC

We purified RRF_mt_ carrying 25 and 79 amino acid (aa) N-terminus extensions (NTEs), and prepared a model PoTC by treating the mitoribosome with puromycin (see Methods). The RRF_mt_ carrying full 79 aa NTE binds efficiently to the model mitoribosome PoTC, whereas the RRF_mt_ construct carrying only 25 aa NTE does not show detectable binding and therefore was not pursued for structural studies. We obtained an overall 3.7 Å resolution structure of the mitoribosome-RRF_mt_ complex (Fig. S1c), which showed a clear density for RRF_mt_, but a relatively weak density for the small (28S) mitoribosomal subunit, suggesting more than one conformational states for the 28S subunit. Therefore, the dataset was further classified to capture conformational states of the 28S subunit.

### RRF_mt_ preferentially binds the rotated 55S mitoribosome

After 3D classification, we obtained four subpopulations from a total of 144,051 selected particle images that included three major classes corresponding to intact 55S mitoribosomes with and without bound RRF_mt_, dissociated large (39S) mitoribosomal subunit (Fig. S1c), and a minor class that contained mostly damaged particles. The 55S classes with and without bound RRF_mt_ were refined to 3.9 Å and 4.4 Å resolution, respectively. Superimposition of the RRF_mt_-bound 55S map (henceforth referred to as Class I) with the published similar resolution cryo-EM maps of the human ^24^ and porcine ^25^ 55S mitoribosomes, both in their classical unrotated conformations, revealed a pronounced rotation of the 28S subunit relative to the 39S subunit in our structure. This rotation is similar to the ratchet-like intersubunit rotation observed in bacterial ribosome, where the small subunit shows a coordinated 5-10° rotation in a counter-clockwise direction with respect to the large subunit ^39,40^, with largest and smallest displacements taking place in the peripheral and central regions, respectively, of the small subunit. In our 55S-RRF_mt_ structure, the 28S subunit rotates by ∼ 8.5° with respect to the 39S subunit (Fig. S2a), using a pivot point of rotation at the 12S rRNA base A1162 within h27 (all rRNA helices are identified according to bacterial numbering ^41,42^) on the 28S body. This movement results in maximum displacement involving the mito-specific protein S22 by ∼15 Å that lies at the bottom of the 28S-subunit body. In addition to the ratchet-like rotation of the whole 28S subunit, the head of the 28S subunit rotates by ∼ 5° in an orthogonal direction towards the E site, which is similar to the swiveling movement described previously ^43,44^ (Fig. S2b), resulting in placement of the 28S head protein S29 further away from the 39S subunit.

Interestingly, the conformation of the empty 55S cryo-EM map (henceforth referred to as Class II) is not only different from our RRF_mt_-bound 55S map, but also from the published bovine ^23^ and porcine ^25^ 55S mitoribosome structures. This conformational change involves a ∼ 8° rotation of the 28S subunit around a long axis of the subunit such that its shoulder side moves towards 39S subunit while its platform side moves away from the 39S subunit. Previous cryo-EM studies of the empty human 55S mitoribosome ^24^ and posttranslocational-state mammalian 80S ribosome ^45^ have also reported the existence of similar conformation that is termed as “subunit rolling” ^45^, but thus far has not been reported for any bacterial ribosomal complex. The fact that the 28S subunit in our empty 55S mitoribosome map (Class II) does not adopt a ratchet-like rotation suggests that a fully-rotated conformational state of the 55S mitoribosome may be a prerequisite for binding of RRF_mt_. It should be noted that the ribosome in eubacterial ribsosome-RRF complex is almost invariably captured in a ratcheted state ^12-16^; however, it is not clear whether RRF binding induces such a state or it binds to a pre-ratcheted ribosome.

Intersubunit bridges between the two subunits of the ribosome are essential for functionally intact ribosome ^23-25,46,47^ that allow dynamic movements required to facilitate various steps of the translation cycle. There are about fifteen intersubunit bridges that have identified in the mammalian 55S ribosome ^23,24^ and seven of them are either destabilized or completely broken during ratchet-like intersubunit rotation ^24^. Interestingly, three (mB1a, mB1b and mB2) out of these seven bridges are specific to the mitoribosomes, involving mito-specific proteins S29, L40, L46 and L48 ^24,25^. It appears that the RRF_mt_ binding stabilizes the rotated conformational state of the mitoribosome with multiple weakened mito-specific intersubunit bridges to prepare the complex for the subsequent step of subunit dissociation upon EF-G2_mt_ binding.

### Molecular Interactions of RRF_mt_ with components of the 55S ribosome

In our cryo-EM map of the 55S-RRF_mt_ complex, a well-resolved “L” shaped density readily attributable to RRF_mt_ is visible in the intersubunit space of the 55S mitoribosome (Fig. 1a & b), matching in the overall size and domain composition to that of bacterial RRF on ^12^-^17^ and off ^9,10^ the 70S ribosome. The long arm of the L-shaped density is represented by three long α-helices that run parallel to each other and constitute the larger domain I, while the smaller domain II is composed of an α/β motif (Fig. 1c). Like in bacterial ribosome-RRF complexes, RRF_mt_ in mitoribosome is located on the cleft of the 39S subunit and is positioned such that its domain I is oriented towards the PTC and overlaps with the binding positions of the A- and P-site tRNAs, while domain II is oriented towards the intersubunit space and is positioned close to protein S12. In its current position, domain I will preclude the binding of both A- and P-site tRNAs in their classical A/A and P/P binding states (Fig. 2), but would allow the binding of P-site tRNA in its P/E hybrid state. Thus, the overall binding position of structurally conserved region of domain I of RRF_mt_ in our structure is almost identical to those of domain I of RRF in the bacterial ^12,14^-^17^ and chloroplast ribosomes ^22^, whereas domain II has been found to exhibit different orientations in each of those bacterial ribosome complexes.

**Fig. 1.**
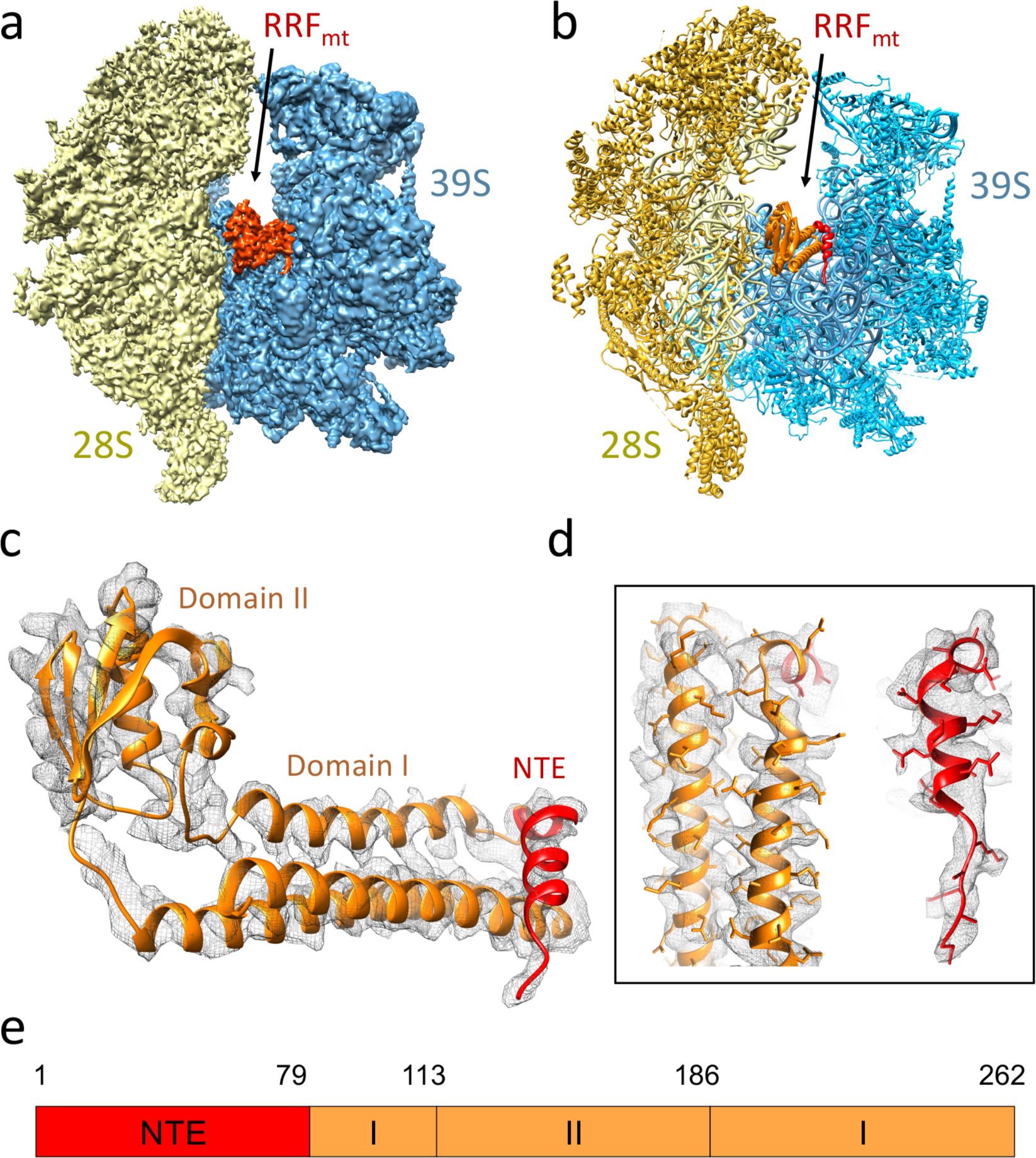
Cryo-EM structure of the human 55S mitochondrial ribosome in complex with RRF_mt_. (**a**) Three-dimensional cryo-EM map of the 55S mitoribosome-RRF_mt_ complex as seen from the subunit-subunit interface side, with segmented densities corresponding to the small subunit (28S, yellow) large subunit (39S, blue), and RRF_mt_ (orange). (**b**) Molecular interpretation of the cryo-EM map shown in panel (a). A darker shade of yellow is used to differentiate the 28S ribosomal proteins from the 12S rRNA, while a lighter shade of blue is used to differentiate the 39S ribosomal proteins from the 16S rRNA. Landmarks of the 28S subunit: h, head; b, body. Landmarks of the 39S subunit: CP, central protuberance. (**c** & **d**) Molecular model of RRF_mt_ as derived from the cryo-EM map, showing well-resolved densities for both the conserved domains I and II (orange) and the NTE (red). The better resolved segments of RRF_mt_ with sidechains are shown in panel (d). (**e**) Overall domain organization of the human RRF_mt_.

**Fig. 2.**
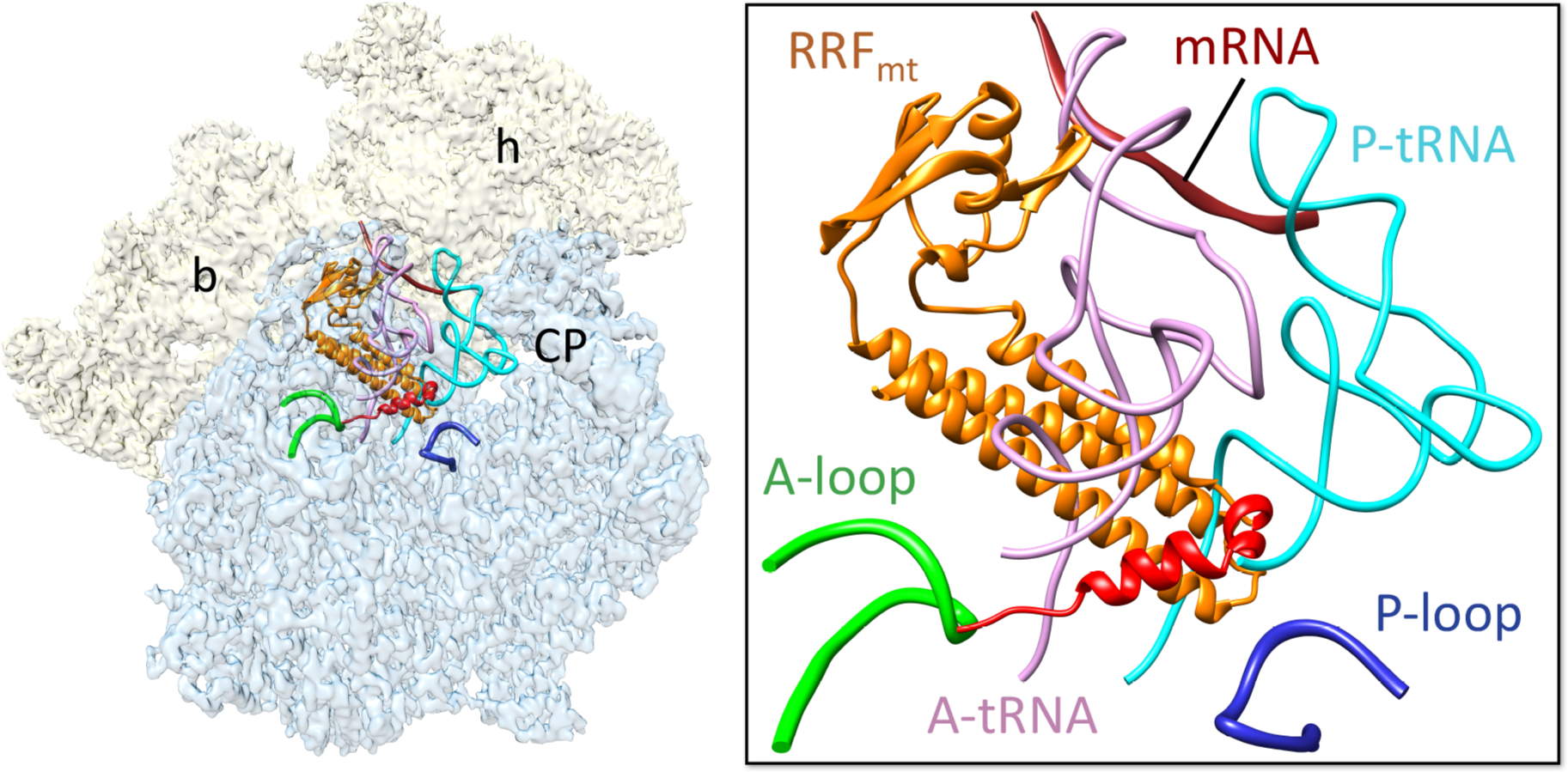
Binding positions of bound RRF_mt_, mRNA, and A- and P-site tRNAs on the mitoribosome. RRF_mt_ binding would be in direct steric clash with the acceptor arm of both aminoacyl- and peptidyl- tRNAs on the large subunit. Coordinates of the mRNA (dark brown) and tRNAs (pink and light blue) were derived from the structure of porcine mitoribosome ^25^ (PDB ID 5AJ4). The conserved RRF_mt_ domains and its NTE are colored as in Fig. 1. The NTE of RRF_mt_ lies in close proximity to the functionally-important and conserved A- (green) and P-loops (dark blue). A thumbnail to left depicts an overall orientation of the 55S mitoribosome, with semitransparent 28S (yellow) and 39S (blue) subunits, and overlaid positions of ligands. Landmarks on the thumbnail: h, head, and b, body of the 28S subunit, and CP, central protuberance of the 39S subunit.

#### Interactions of Domain I

Though the overall shape and composition are significantly different among the bacterial, mitochondrial and chloroplast ribosomes, their internal rRNA cores are very much conserved ^21^-^26^ The strategic positioning of domain I on the 55S mitoribosome allows RRF_mt_ to interact with several important elements of the large subunit, such as components of the PTC, helix 69 (H69) and helix 71 (H71), that are known to play crucial roles during various phases of the bacterial translational cycle ^48^. The tip of domain I is positioned very close to the P-loop region (H80) of the PTC such that its aa residues Ser227-Asp229 make direct contacts with the rRNA bases G2816-G2819 (Fig. 3a). The P-loop region is known to interact with the CCA-end of the P-site bound tRNA, and hence it is essential that the peptidyl tRNA is be removed from the P site of the 39S subunit prior to the binding of RRF_mt_. Interestingly, these interacting residues are conserved among bacterial, mitochondrial and chloroplast RRFs; accordingly, structural studies of the bacterial and chloroplast 70S-RRF complexes ^12-17,21^ have shown similar interactions of domain I with the P-loop. The three long α-helices of domain I run almost parallel to a portion of 16S rRNA helix H71 such that aa residues Asn202, Lys205, Arg209, Arg212, Thr213 and Met216 from α-helix 3 of RRF_mt_ make extensive interactions with the H71 rRNA bases A2604-U2609 (Fig 3b). Surprisingly, except for the two arginine residues (Arg209 and Arg 212), none of the interacting α-helix 3 residues are conserved between the bacterial and the human mitochondrial RRF, and mutation of Arg209 (Arg132 in *E. coli*) to other aa residues drastically reduces the affinity of RRF to the 70S ribosome ^49^. This high network of interactions seems to provide the essential anchoring points for RRF_mt_ on the 39S subunit, as mutations in the analogous region of the prokaryotic factor can lead to cell death or generation of temperature-sensitive phenotypes in bacteria ^50^. Though the α-helix 3 within domain I of RRF_mt_ is positioned in the close vicinity of H69, no specific interactions between domain I and H69 were detected. Interestingly, the presence of RRF_mt_ appears to have no impact on the conformation of H69, a key 16S rRNA helix that partners with the functionally important 12S rRNA helix, h44, to form the highly-conserved and crucial intersubunit bridge B2a in the 55S ribosome. This observation is in sharp contrast to multiple contacts reported between domain I of RRF and H69 in bacterial ribosome-RRF complexes, where H69 has been shown to adopt different conformations upon RRF binding ^12,14,15,18^.

**Fig. 3.**
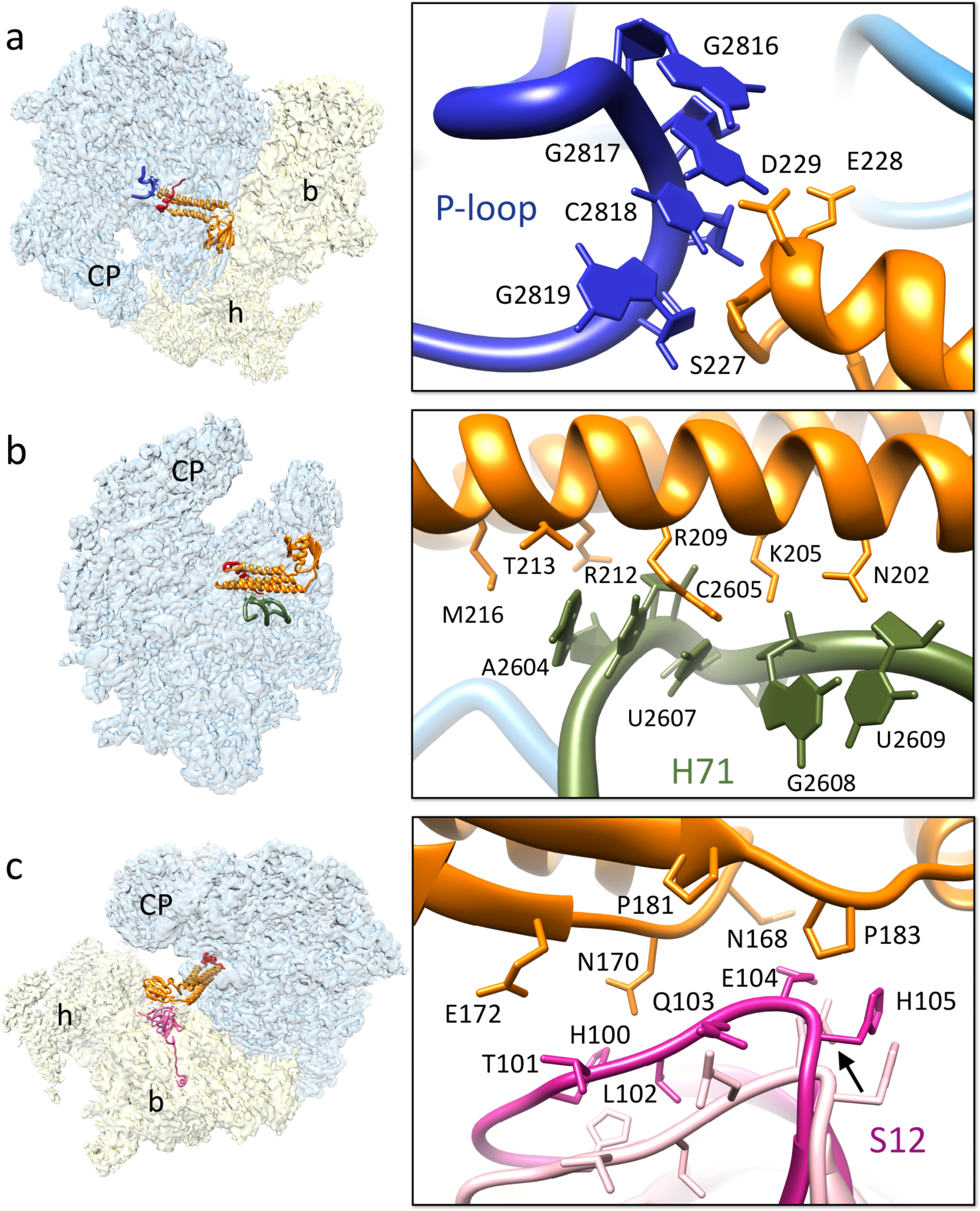
Interactions of structurally conserved domains of RRF_mt_ with the 55S mitoribosome. (**a**) Contacts between the domain I (orange) and the P-loop, the 16S rRNA H80 (blue). (**b**) Interactions between the domain I and H71 (olive green) of the 12S rRNA. (**c**) Contacts between the ribosomal protein S12 (magenta) and RRF_mt_ domain II. Arrow points to a shift in the interacting S12 loop in our structure, as compared to its position (pink) in an unratcheted conformation ^24^ (PDB ID 3J9M). Thumbnails to left depict overall orientations of the 55S mitoribosome, with semitransparent 28S (yellow) and 39S (blue) subunits, and overlaid positions of ligands. Landmarks on the thumbnails are same as introduced in Fig. 2.

#### Interactions of Domain II

The positioning of domain II in the present structure is different as compared to the orientation of domain II in the 70S-RRF ^17^ and the 70S-chlRRF ^21^ complexes. While two RRF domains are placed in almost right angles to each other in most bacterial complexes, domain II adopts a more open conformation in the chloroplast complex. In our structure, domain II occupies an intermediate position relative to the positions of domain II in the bacterial and chloroplast complexes (Fig. S3). None of the ribosomal components from the 39S subunit are found within interacting distance from domain II of RRF_mt_. The only structural element from the 28S subunit that is found in close proximity to domain II of RRF_mt_ is protein S12. A loop region of protein S12 encompassing His100-His105 is within hydrogen-bonding distance (∼ 4 Å) from the asparagine and proline-rich segment that involves aa residues Asn168, Asn170, Glu172, Pro181 and Pro183 of the RRF_mt_ domain II (Fig. 3c). Both the prolines and Asn170 are conserved among the bacterial and human mitochondrial RRFs. Though interactions between S12 and bacterial RRF are proposed to exist in the 70S-RRF structures ^12,14,16^, these interactions are more extensive in our 55S-RRF_mt_ complex. Superimposition of the published human 55S ribosome map ^24^ with our Class I map reveals large conformational change in S12, where His100-His105 loop is shifted by ∼5 Å towards the RRF_mt_ domain II (Fig. 3c). This observation implies that either the conformation of S12 in the rotated 28S subunit stabilizes the binding of RRF_mt_ domain II, or the interaction between S12 and RRF_mt_ domain II induces a local conformational change that could lead to subunit rotation.

#### Interactions of the mito-specific N-terminal extension (NTE)

The human RRF_mt_ carries a functionally essential mito-specific 80 amino acid extension at its N-terminus ^37,38^ (Fig. S3). In our map, we find additional density (Fig. 1c, d) at the apical region of RRF_mt_ domain I that is contiguous with the RRF_mt_ α-helix1 and could be readily assigned to the last 21 aa residues of the NTE. Residues Lys65-Val79 fold into a well-defined α-helix, residues Lys59-Gly64 constitute a loop region (Fig. 1c, d), while the remaining residues of the NTE could not be modelled due to lack of a consistent density. The observed NTE segment runs beyond α-helix 4 in an oblique perpendicular direction to rest of the domain I, and is found sandwiched between the 39S subunit and rest of the RRF_mt_ domain I, where it makes several novel interactions with the 39S subunit.

The ribosomal elements that are within reach of the NTE are 16S rRNA helices H89, H90 and H92, and ribosomal proteins L16 and L27. Protein L27 lies in close proximity to the PTC and was reported to interact with the apex region of domain I in the bacterial recycling complex ^15,17,18^. The N-terminal region of L27 is known to be highly flexible and hence was disordered in the factor-free ^44,46^ as well as RRF-bound 70S ribosomal complexes ^17^. Human mitoribosomal protein L27 is made up of 148 aa residues, of which the first 30 residues constitute the mitochondrial target sequence (MTS) that would be eventually cleaved off in the mature protein. Thus, the mature L27 would be ∼118 aa, which is still larger than its bacterial counterpart. In our structure, we have observed additional density extending from the previously known N-terminus of L27, as compared to that in empty human 55S ribosome structure ^24^, suggesting that contacts with the NTE of RRF_mt_ has stabilized the N-terminus of the L27 in our map, and therefore, allowed us to model additional aa residues, except for the first three amino acid residues. As compared to the empty human and porcine 55S ribosomal structures ^24,25^, the N-terminal region of L27 has undergone a large conformational change, and moves by ∼9 Å towards the P-site tRNA (Fig. 4a) such that it would occlude the binding of the accepter arm of the peptidyl-tRNA on the 39S subunit (Fig. 4a). The N-terminus of L27 makes multiple contacts with the mito-specific NTE of RRF_mt_ through its residues Asn35 and Lys34. Asn35 interacts with Ala77 and Leu78, whereas the conserved Lys34 makes close contacts with Asn75 and Lue78 from the NTE of RRF_mt_ (Fig. 4b). By interacting with the CCA end of the P-site bound tRNA and thereby stabilizing it, L27 was believed to enhance the peptidyl transferase activity at the PTC ^46,51^. In a similar fashion, L27 is positioned to stabilize the binding of RRF_mt_ through the novel mito-specific interactions described here. The large subunit protein L16 is known to be in close proximity to bacterial RRF ^14,17,18^, but specific interactions between the two proteins could not be identified in previous studies. L16 and the NTE of RRF_mt_ contact each other at multiple sites. Residues Ser135-His138, Ala146 and Asp148 from the L16 are involved in direct interactions with the NTE residues Thr70, Lys71, Ile74 and Asn75 (Fig. 4c). Outside the NTE, Leu233 from RRF_mt_ domain I also makes a close contact with His138 of L16. L16 in human mitochondria has high clinical significance as it gets upregulated in breast cancer ^52^ and colorectal cancer ^53^. Direct interactions observed between RRF_mt_ NTE and L16 could provide new insights in to role of L16 in these disease conditions.

**Fig. 4.**
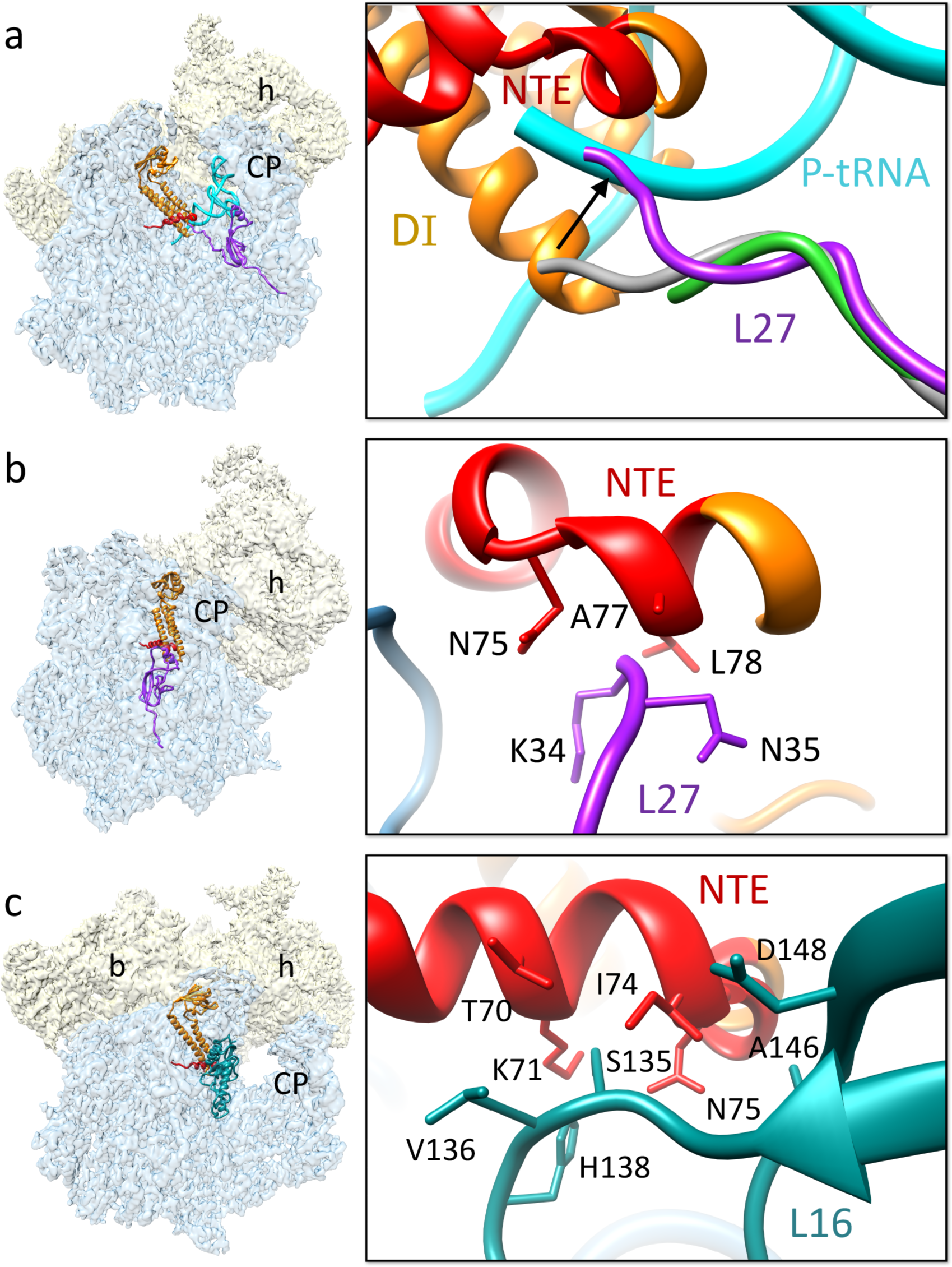
Mito-specific Interactions between the NTE of RRFmt NTE and the 39S mitoribosomal subunit proteins. (**a**) The N-terminal region of L27 (purple) in the human 55S-RRF_mt_ complex has shifted by ∼9 Å towards the peptidyl tRNA-binding site as compared to the positions of L27 in the porcine (grey) ^25^ (PDB ID 5AJ4) and empty human (green) ^24^ (PDB ID 3J9M) mitoribosomes to interact with the NTE (red) of RRF_mt_. In its new conformation, L27 would also block the binding of tRNA in the 39S P site. Mito-specific interactions between the NTE and mitoribosomal proteins are shown in panels (**b**) L27 and (**c**) L16 (dark cyan). Thumbnails to the left depict overall orientations of the 55S mitoribosome, with semitransparent 28S (yellow) and 39S (blue) subunits, and overlaid positions of ligands. Landmarks on the thumbnails are same as introduced in Fig. 2.

The RRF_mt_ interaction with the 16S rRNA helix 92 (H92) is of particular interest because it has the functionally important A-loop that is known to interact with the CCA end of the A-site tRNA, and is thus associated with PTC (Fig. 5a). Lys61, from the RRF_mt_ NTE interacts with nucleotides C3043 and A3044 of the 16S rRNA H92, while the neighboring Ala60 also contacts C3043 (Fig. 5b). It should be noted that this stretch of the NTE has unusual presence of multiple lysine residues. RRF_mt_ NTE residues Ala62 and Lys63 interact with the 16S rRNA H89 base U2979 (Fig. 5b), and residues Gly64 and Gln67 is positioned very close to the neighboring base U2980. It should be noted that H89 extends up to the GTPase-associated center (GAC) of the L7/L12 stalk base ^47^, and its interactions with NTE might influence the subsequent EF-G2_mt_- catalyzed GTP hydrolysis step. Unlike the 70S ribosomal recycling, GTP hydrolysis on EF-G2_mt_ is not required for the splitting of the 55S ribosome, and the hydrolysis reaction is only necessary for the release of EF-G2_mt_ from the mitoribosome ^36^. A strategically placed lysine residue Lys59 from the NTE interacts with the backbone phosphates of bases G3054 and U3055 on the 16S rRNA H90 through charge-based interactions (Fig 5b). Interestingly, all three rRNA helices are part of the domain V of 23S rRNA that comprise the highly-conserved PTC (Fig. 5), while helices H89-93 constitute the region of domain V that is known for stabilizing the elongation factor-binding region of the 70S ribosome ^41^. Simultaneous interactions of the RRF_mt_ NTE with the three 16S rRNA helices that have been implicated in EF-G binding (H90, H92) and GTP hydrolysis (H89) would have direct effect on the subsequent EF-G2-mediated steps during the mitoribosome recycling process.

**Fig. 5.**
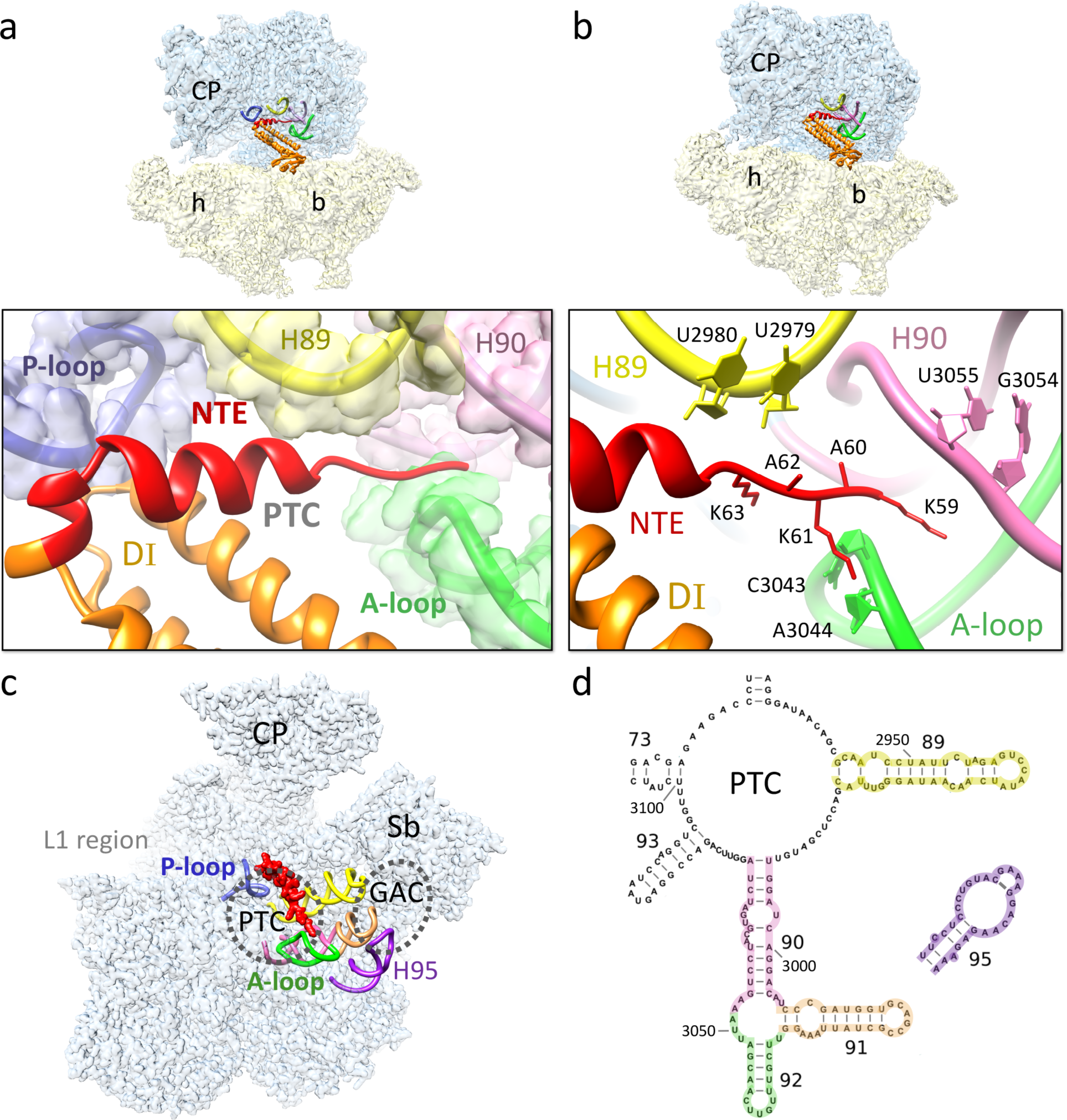
Mito-specific interactions between the NTE of RRF_mt_ and the 16S rRNA components of the 39S subunit. (**a**) The NTE (red) of RRF_mt_ is strategically positioned in the peptidyl-transferase center (PTC) to interact with several functionally important 16S rRNA segments such as the P-loop (blue), H89 (yellow), H90 (pink) and the A-loop (H92) (green). (**b**) Mito-specific interactions of the NTE with the nucleotide bases of H89, H90 and the A-loop. Color codes are same as in (a). Thumbnails on the top depict overall orientations of the 55S mitoribosome in panels a and b, with semitransparent 28S (yellow) and 39S (blue) subunits, and overlaid positions of ligands. Landmarks on the thumbnails are same as introduced in Fig. 2. (**c**) Overall location of the NTE on the 39S subunit (semitransparent blue) as seen from the subunit’s interface side, with the rRNA components of PTC, including extended rRNA helices that run up to the GTPase-associated center (GAC), which includes H95 that carries the α-sarcin-ricin stem loop. Both PTC and GAC are depicted as dashed circles. (**d**) Secondary structure of the PTC region of the 16S rRNA, highlighting helices that are color coded as in panels a and c.

In conclusion, our study provides the first direct visualization and structure of the human mitoribosome-bound human RRF_mt_ and its mito-specific NTE, and their interactions with the functionally relevant components of the mitoribosome. These include both A and P loops of the PTC ^20^, and components of the PTC that directly communicate with the GTPase-associated center of the large ribosomal subunit. Our observation that the mito-specific RRF_mt_ NTE spans across A and P loops, encompassing the entrance of the nascent polypeptide-exit tunnel, suggests that NTE interaction with the mitoribosome ensures a complete inaccessibility to tRNAs and other ligands to both PTC and the entrance nascent polypeptide-exit tunnel of the mitoribosome. This situation is in sharp contrast to that in bacterial RRF, which directly interacts only with the P loop. Extensive interactions between the RRF_mt_ and mitoribosome is a novel feature of RRF_mt_-mediated mitoribosome recycling. Future structural studies involving EF-G2_mt_ should provide further insights into the precise functional roles of the new interactions observed in the present study.

## Methods

### Isolation of mitochondria from HEK cells

The source of mitochondrial ribosomes was human embryonic kidney cells lacking N-acetyl-glucosaminyltransferase I (HEK293S GnTI) that were cultured in roller bottles using FreeStyle^TM^293 media (Gibco, Life Technologies) supplemented with 5% fetal bovine serum (Gibco, Life Technologies). After centrifugation at 1,000 x g for 7 minutes, the HEK293S GnTI cell-pellet was transferred to a glass homogenizer and resuspended in buffer containing 50 mM HEPES-KOH pH 7.5, 10 mM KCL, 1.5 mM MgOAc, 70 mM sucrose, 210 mM mannitol, 1 mM EDTA, 1 mM EGTA, 1 mM DTT, and 1 mM PMSF. The cells were homogenized by applying 120 strokes and the supernatant was separated from the cell debris by spinning at 950 x g for 15 minutes. The supernatant was then spun at 11,000 x g for 15 minutes, and the resulting pellet that contains crude mitochondria was resuspened in SEM buffer (250 mM sucrose, 20 mM HEPES-KOH pH 7.5, 1 mM EDTA, and 1 mM EGTA) followed by DNase treatment. A discontinues sucrose gradient was prepared in buffer containing 10 mM HEPES-KOH pH 7.5 and1 mM EDTA) and the DNase-treated sample was loaded over the gradient and centrifuged for one hour at 135,000 x g using Ti70 rotor in Beckman ultracentrifuge. The brownish-orange layer containing pure mitochondria was carefully separated and re-suspended in SEM buffer. The sample was then spun at 10,000 x g for 15 minutes, and the pellet was stored at −80°C.

### Isolation of mitoribosomes from mitochondria

Four volumes of lysis buffer (25 mM HEPES-KOH pH 7.5, 100 mM KCl, 25 mM MgOAc, 1.7 % Triton X-100, 2 mM DTT and 1 mM PMSF) was added to the mitochondrial-pellet and then incubated for 15 minutes at 4°C. The sample was centrifuged at 30,000 x g for 20 minutes and the supernatant was loaded on top of 1 M sucrose cushion in buffer (20 mM HEPES-KOH pH 7.5, 100 mM KCl, 20 mM MgOAc, 1% Triton X-100 and 2 mM DTT). After centrifugation for 17 hours at 90,000 x g using Ti70 rotor in Beckman ultracentrifuge, a minimal volume of Mitobuffer (20 mM HEPES-KOH pH 7.5, 100 mM KCl, 20 mM MgOAc, and 2 mM DTT) enough to dissolve the pellet was added. A continuous sucrose gradient (10-30%) was prepared in Mitobuffer, the resuspended pellet was subjected to density gradient centrifugation at 60,000 x g for 17 hours using Sw32 rotor in Beckman ultracentrifuge. The gradient was fractionated on ISCO gradient analyzer (Teledyne ISCO, Inc), and the fractions corresponding to the mitoribosomes were collected and pooled. Finally, the pooled mitoribosomes were concentrated by spinning them at 130,000 x g for 6 hours using Ti70 rotor, and the pellet was resuspended in Polymix buffer (5 mM HEPES-KOH pH 7.5, 100 mM KCl, 20 mM MgOAc, 5 mM NH_4_Cl, 0.5 mM CaCl_2_, 1 mM DTT, 1 mM spermidine, and 8 mM putrescine).

### Purification of RRF_mt_

RRF_mt_ clone was a gift from Prof. Linda Spremulli, University of North Carolina. The SUMO-tagged RRF_mt_ was over-expressed in Rosetta2 cell lines and lysis buffer (500 mM NaCl, 1X PBS, 4 mM ß-mercaptoethanol, 1 mM PMSF, and 10 mM imidazole) was added to the pelleted cells. After sonication, the lysate was treated with DNase and then centrifuged for 30 minutes at 16,000 x g. The supernatant was applied to a His-trap Ni^2+^ column and the SUMO-tagged protein was eluted from the column using elution buffer (250 mM NaCl, 1X PBS, 4 mM ß-mercaptoethanol, and 300 mM imidazole) using standard affinity purification protocols. The purified protein was dialyzed in buffer (20 mM Tris-HCl pH 8.0, 250 mM NaCl, 4 mM ß-mercaptoethanol, and 5% glycerol) and then the SUMO tag was cleaved from RRF_mt_ by incubation with SUMO express protease for 1 hour at 30°C. Finally, the SUMO tag and the SUMO express protease were separated from the RRF_mt_ by passing through a His-trap Ni^2+^ column that specifically adsorbs the SUMO tag and the SUMO express protease while pure RRF_mt_ was released into the column flow through.

### Preparation of the 55S-RRF_mt_ complex

50 mM puromycin was added to 100 nM 55S mitochondrial ribosomes in Polymix buffer and incubated for 10 minutes at 37°C to release any P/P-state tRNAs from the large mitoribosomal subunit. This step was necessary to free up the anticipated RRFmt binding site on the mitoribosome. This step was followed by the addition of 10 µM RRF_mt_ and an additional 5 minutes incubation at 37°C. The reaction mixture was then immediately utilized for grid preparation for cryo-EM analysis.

### Cryo-Electron microscopy

Quantifoil holey copper 1.2/1.3 grids were pre-coated with a thin layer (∼ 50 Å thick) of home-made continuous carbon film and glow-discharged for 30 seconds using a plasma sterilizer. 4 µl of the sample was applied to the grids, incubated for 15 seconds at 4°C and 100% humidity and then blotted for 4 seconds before flash-freezing into the liquid ethane using Vitrobot (FEI company). Data was acquired on a Titan Krios electron microscope (FEI company) equipped with a Gatan K2 summit direct-electron detecting camera at 300 KV. A defocus range of 0.8 to 2.5 µm was used at a calibrated magnification of 105,000 X, yielding a pixel size of 1.09 Å on the object scale. A dose rate of 7 electrons per pixel per second and an exposure time of 10 seconds resulted in a total dose of 70 eÅ^-2^. After determining their contrast transfer function (CTF) parameters using CTFFIND4 ^54^, the bad micrographs were deselected from the good ones. The data was further processed in Relion 2.0 ^55^ and a total of 288,138 particles were picked from the selected 5,305 micrographs using the auto-pick function. CryoSPARC ^56^ was used to perform all the subsequent 2D and 3D classifications and refinements. From 288,138 autopicked particles, 144,057 good particles were selected after reference-free 2D classification. Particles in the selected 2D averages were then subjected to an initial reference-based heterogeneous 3D classification that yielded four major 3D classes (Fig. S1). The Class I and the Class II turned out to be 55S ribosomes in their ratcheted and unratcheted conformational states, respectively. The Class I that contained 67,116 particles showed a strong density for the ligand RRF_mt_, whereas the Class II contained 26,195 particles and did not have any density for RRFmt. The Class III that contained 42,352 particles was purely made up of 39S mitoribosomal subunits, while the Class IV with 8,394 particles was regarded as junk since it did not produce any defined structure. Class I, Class II and Class III were further refined to 3.9 Å, 4.4 Å and 4.3 Å resolution, respectively.

### Model building and refinement

The model of the 55S mitoribosome was generated by using the previously published coordinates of human mitoribosome ^24^ (PDB ID: 3J9M) as a template. Coordinates of the 28S and 39S subunits were docked independently as rigid bodies into the corresponding cryo-EM densities using Chimera ^57^. The models were further real-space refined and validated in PHENIX ^58^ to obtain optimal fitting to our cryo-EM densities. The amino acid sequence of the human RRF_mt_ was obtained from the NCBI protein data base and the sequence was submitted to Robetta server ^59^ for initial homology model prediction. Robetta generated five homology models. All five models were placed into the corresponding cryo-EM density as rigid bodies using Chimera 1.11 ^57^ and the model that has the best matching features with the cryo-EM density was selected. The crystal structure of 70S bound *T. thermophilus* RRF ^17^ (PDB ID: 4V5A) was used as reference and regions in the RRF_mt_ that are not fully accommodated into the cryo-EM density were modelled in Chimera 1.11 ^57^. Human RRF_mt_ contains a long N-terminal extension (NTE) which was built *de novo* using the “Build structure” function in Chimera 1.1 ^57^. The NTE was modelled based on the recognizable secondary structural elements and the position of bulky side chains in the cryo-EM density. Finally, we used the “Real-space refinement” function in PHENIX for optimization of the model into the cryo-EM density and further validation of the model ^58^.

## Acknowledgements

Authors acknowledge use of the Wadsworth Center’s Media and Culture core facility for help in producing large volumes of HEK293S GnTI cells. We acknowledge the use of the Wadsworth Center’s and New York Structural Biology Center’s (NYSBC’s) EM facilities. NYSBC EM facilities are supported by grants from the Simons Foundation (349247), NYSTAR, the NIH (GM103310) and the Agouron Institute (F00316). We thank Mona Gupta and Prem Kaushal for their initial contributions to the project, ArDean Leith for help with computation, and Nilesh Banavali for critical reading of the manuscript. This work was supported by the NIH grant R01 GM61576 (to R.K.A.).

## Author Contributions

RKA conceived this study. RKK and PK performed biochemical experiments. PR collected cryo-EM data. RKK, and MRS performed image processing. RKK, MRS, and RKA analyzed the data and wrote the manuscript. Authors declare no competing financial interest in this work.

## Data Deposition

The cryo-EM maps and atomic coordinates of the 55S mitoribosome-RRF_mt_ complex has been deposited in the Electron Microscopy and PDB Data Bank (wwPDB.org) under accession codes EMD----- and PDB ID, respectively.

